# Maintaining proton homeostasis is an essential role of glucose metabolism in cell survival

**DOI:** 10.1101/058594

**Authors:** Yanfen Cui, Yuanyuan Wang, Pan Xing, Li Qiu, Miao Liu, Xin Wang, Guoguang Ying, Binghui Li

## Abstract

Aerobic glycolysis, termed “the Warburg Effect”, supports cell proliferation, and glucose deprivation directly elicits necrosis or shifts stimuli-induced apoptosis to necrosis. However, how glucose metabolism regulates cell survival or death choice remains largely unclear. Here we use our recently developed method to monitor in real-time cellular apoptosis and necrosis, and uncover a metabolic homeostasis linked to cell death control. We show that glucose metabolism is the major source to maintain both intracellular and extracellular proton homeostasis. Glucose deficiency leads to lack of proton provision, which provokes a compensatory lysosomal proton efflux and resultant increased lysosomal pH. This lysosomal alkalinization can trigger necrosis. Furthermore, artificial proton supplement enables cells to survive glucose deprivation. Taken together, our results reveal a critical role of glucose metabolism in maintaining cellular microenvironment, and provide a better understanding of the essential requirement of aerobic glycolysis for proliferating cells whose active anabolism consumes a great many protons.

## Introduction

Unlike non-proliferating cells, which rely on mitochondrial oxidative phosphorylation to generate ATP, proliferating cells, in particular cancer cells, are apt to rely on aerobic glycolysis, an inefficient way to generate energy (Vander Heiden et al., 2009). Since 1920s when this phenomenon, termed “the Warburg Effect”, was observed (Warburg, 1956), its growth advantage provided for proliferating cells has been largely unknown. Although other metabolites, such as pyruvate and amino acids, can efficiently generate ATP in mitochondrial pathway or can even be converted to glucose through gluconeogenic pathway (Zhang et al., 2014), glucose deficiency eventually leads to cell death (Yun et al., 2009). These observations suggest that glucose catabolism plunks for cell survival independently of ATP generation.

In addition, glucose deprivation has been known as a factor that switches stimuli-induced apoptosis to necrosis in some cell types (Eguchi et al., 1997; Leist et al., 1997). Although the detailed mechanism need to be further dissected, glucose metabolism seems to be certainly involved in determining how cell death occurs. Metabolism, in essence, is a pool of self-sustaining chemical transformations within the cells, and these reactions function as the fundament of living cells and allow cells to survive and grow. Perturbations of metabolic homeostasis by direct assaults or by signal cascades often lead to apoptosis or necrosis. Therefore, like cell survival, cell death should also be under tight metabolic control (Green et al., 2014). Over the past few decades, most of the effort has been focused on establishing molecular connections between signal transduction pathways and cell death. Recently, it has become clear that many signaling pathways converge to regulate metabolism to support cellular processes, including cell death. However, how metabolism governs cell survival, proliferation or death remains largely unknown.

Due to deficiency in caspase-3, MCF-7 cells exposed to TNF-α undergo caspase-dependent apoptosis through a well-characterized pathway but do not form apoptotic bodies (Janicke et al., 1998). Moreover, the dead MCF-7 cells do not detach from the culture platform. Therefore, the MCF-7 cell line is an ideal model for fluorescence microscopy-based quantification of cell death. Taking advantage of our recently developed caspase activity reporter that indicates apoptosis by green fluorescence, and using propidium iodide in culture medium to label necrotic cells with red fluorescence (Zhang et al., 2013), we study the effect of metabolism on TNF-α-induced cell death in MCF-7 cells in various nutrient-defined media. We show that cell death and survival are tightly controlled by glucose metabolism-maintained proton homeostasis together with mitochondria and lysosomes.

## Results

### TNF-α-induced necrosis in MCF-7 cells depends on glucose deprivation

We recently developed an sfGFP-based caspase-3-like protease activation indicator (GC3AI) that acquires fluorescent activity only after cleavage by active caspase-3-like proteases during apoptosis (Zhang et al., 2013). In this study, GC3AI was stably expressed in MCF-7 cells to monitor apoptosis. Upon TNF-α treatment for 24h, most of MCF-7/GC3AI cells produced robust green fluorescence with condensed chromatin and shrunken morphology (Figures 1A and 1B), indicating TNF-α-induced apoptosis, as we previously reported (Zhang et al., 2013). In contrast, TNF-α failed to induce green fluorescence and the typical shrunken morphology when MCF-7 cells were deprived of glucose. Hoechst staining showed that chromatin was largely degraded (Figure 1A). Furthermore, time-lapse imaging clearly showed that MCF-7 cell membrane was ruptured and the intracellular contents were released as indicated by Venus fluorescent protein (Figure 1C). This suggests that necrotic cell death is induced by TNF-α plus glucose deprivation.

**Figure 1.**
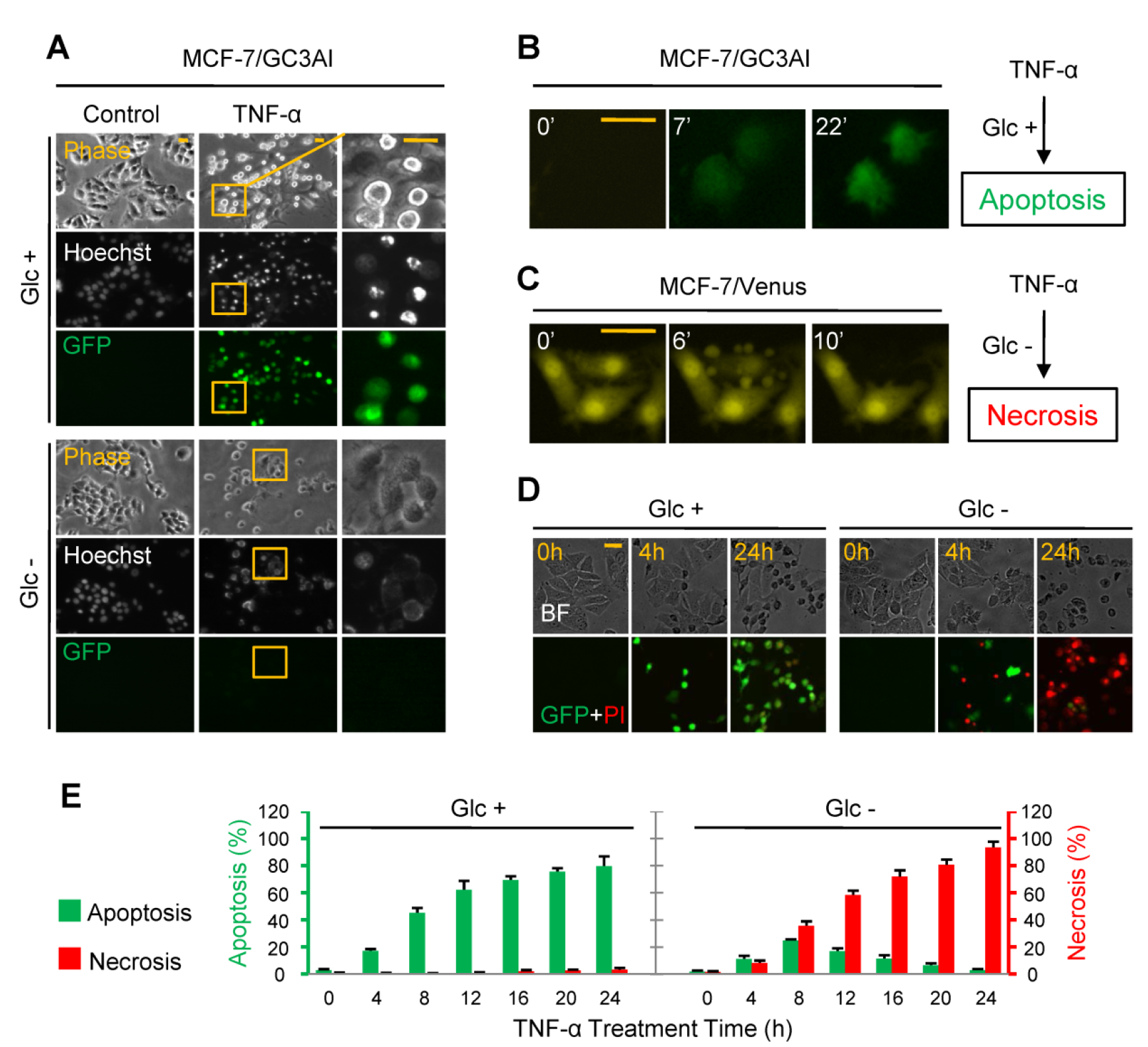
TNF-μ induces apoptosis in the presence of glucose but necrosis in the absence of glucose. (A) Fluorescent images of the fixed MCF-7/GC3AI cells after treatment with 50 ng/ml of TNF-μ for 24h in the presence or absence of glucose and staining with Hoechst. The boxed regions are expanded on the right. (B) Time-lapse images of MCF-7/GC3AI cells after treatment with 50 ng/ml of TNF-μ for 4h in the presence of glucose. GFP fluorescence indicates the activated GC3AI. (C) Time-lapse images of MCF-7/Venus cells after treatment with ng/ml of TNF-μ for 4h in the absence of glucose. Venus fluorescence indicates the cellular morphology. (D) TNF-μ induces apoptosis in the presence of glucose while necrosis in the absence of glucose. GFP fluorescence indicates apoptosis, and PI red fluorescence indicates necrosis. The culture medium contained 2 μg/ml of PI. MCF-7/GC3AI cells were treated with ng/ml of TNF-μ. (E) The fraction of apoptotic and necrotic cells after MCF-7/GC3AI cells were treated with ng/ml of TNF-μ for different times as indicated in the presence or absence of glucose. Error bars indicate ± SD (n=3). Scale bar in all panels, 20 μm. See also Figure S1.

To conveniently monitor in real-time apoptotic and necrotic cell death simultaneously in MCF-7/GC3AI cells, 2 μg/ml of propidium iodide (PI) was added to the culture medium to stain necrotic cells. In the presence of glucose, all TNF-α-induced apoptotic cells displayed green fluorescence at least up to 24h (Figure 1D). By contrast, many PI-positive cells without green fluorescence, referred to as necrotic cells, were observed after TNF-α treatment in the absence of glucose (Figure 1D). Without TNF-α treatment, glucose deprivation alone induced cell death in about 15% of MCF-7 cells at 24h, but showed no effect at 8h (Figure S1A). Interestingly, our time-course results showed that TNF-α also induced apoptosis in addition to necrosis in the absence of glucose starting at 4h (Figures 1D and 1E). However, TNF-α-induced apoptotic cells disappeared by 24h of treatment (Figure 1E), suggesting that at least some necrotic cells were shifted from apoptosis in the absence of glucose. Indeed, such a death transition with morphological change was observed (Figure S1B).

Our results further indicated that the withdrawal of glutamine, pyruvate, amino acids or serum showed no effect on TNF-α-induced apoptosis (Figure S1C). As the withdrawal of glucose as well as other nutrients did not significantly affect the intracellular ATP contents (Figure S1D), TNF-α-induced necrosis in MCF-7 cells was most likely only related to glucose deprivation and independent of ATP depletion. Indeed, a low concentration(>0.1g/l) of glucose was enough to support TNF-α-induced apoptosis at 6h of treatment (Figures S1E and S1F). Taken together, our data indicate that the same apoptotic stimulus, TNF-α, could trigger apoptosis or necrosis in MCF-7 cells depending on the presence or absence of glucose, consistent with the previous observations in other cell lines(Eguchi et al., 1997; Leist et al.,1997).

### Mitochondrial impairment promotes glucose deprivation-induced necrosis

Since MCF-7 cells are deficient in caspase-3, they underwent mitochondria-dependent apoptosis upon TNF-α treatment (Figure 2A)(Scaffidi et al.,1998). This was confirmed by our results that the blockade of caspases by their inhibitors or shRNAs and the over-expression of Bcl-2/Blc-xL almost completely suppressed apoptosis in the presence of glucose (Figures 2B –2D and S2A-S2C). In the absence of glucose, a pan-caspase inhibitor, z-VAD, and caspsase-8 inhibitor, z-IETD, completely blocked TNF-α-induced cell death in MCF-7 cells, whereas the caspase-7 and caspase-9 inhibitors, z-DEVD and z-LEHD, only repressed apoptosis while leaving necrosis unaffected (Figures 2B, 2C and S2A). Similar results were obtained with knockdown of these caspases (Figure S2B). These observations indicate that TNF-α-induced necrosis can occur independently of apoptosis in the condition of glucose deprivation. Since Bcl-2 and Bcl-xL over-expression showed the similar protective effects against TNF-α-induced cell death regardless of glucose (Figure 2D), the two forms of cell death seems to share the upstream pathway and diverge at the mitochondrion (Figure 2A).

**Figure 2.**
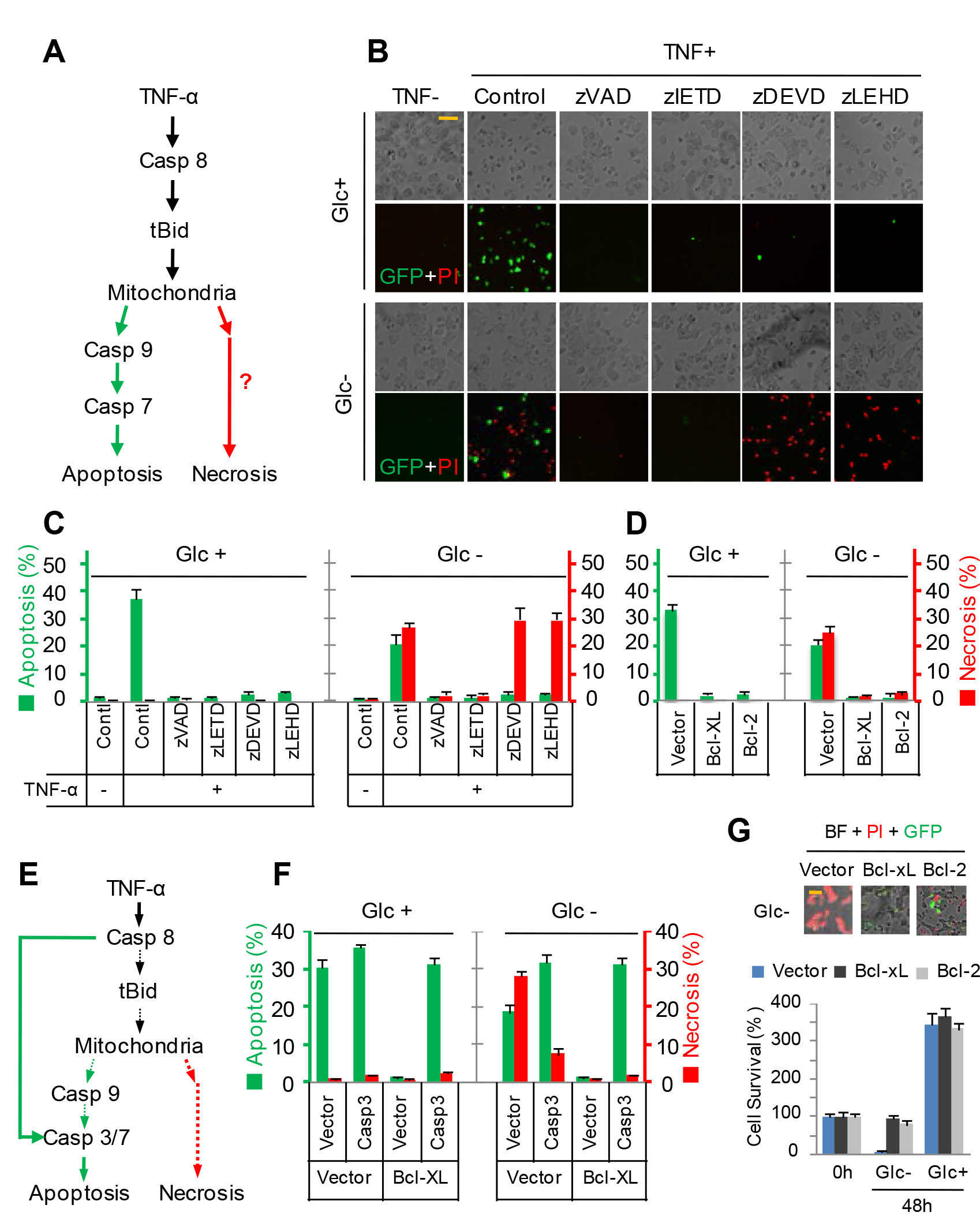
Necrosis induced by TNF-α in the absence of glucose depends on mitochondria. (A) TNF-α-induced mitochondria-dependent apoptotic pathway in MCF-7 cells. (B) Fluorescent images of MCF-7/GC3AI cells after treatment with 50 ng/ml of TNF-α for 6h in the presence or absence of glucose. Cells were pre-treated with 50 μM of z-VAD, z-IETD, z-DEVD, or z-LEHD. Scale bar, 40 μm. (C) The fraction of apoptotic and necrotic MCF-7/GC3AI cells after treatments as described as in (B). (D) The fraction of apoptotic and necrotic MCF-7/GC3AI cells stably expressing Bcl-2 or Bcl-xL. Cells were treated with 50 ng/ml of TNF-α for 6h in the presence or absence of glucose. (E) TNF-α-induced apoptotic pathway in MCF-7 cells stably expressing caspase-3. (F) The fraction of apoptotic and necrotic MCF-7/GC3AI cells stably expressing Bcl-xL and/or caspase-3. Cells were treated with 50 ng/ml of TNF-α for 6h in the presence or absence of glucose. Error bars in all panels indicate ± SD (n=3). (G) Survival of MCF-7/GC3AI cells stably expressing empty vector, Bcl-2 or Bcl-xL in the complete medium or glucose-free medium. Error bars indicate ± SD (n=3). See also Figure S2 and S3.

Bid knockdown partially inhibited TNF-Bid knockdown partially inhibited TNF-induced apoptosis in the presence or absence of glucose while almost completely prevented necrosis in MCF-7 cells in the absence of glucose (Figure S3A). Notably, in MCF-7 cells with Bid knockdown, TNF-α induced the similar level of apoptosis regardless of glucose. Similar effects by TNF-α were also observed in MDA-MB-231 and Bcap37 cells where TNF-α plus cycloheximide induced mitochondria-independent apoptosis (Figures S3B and S3C). Therefore, Bid knockdown may force MCF-7 cells to undergo mitochondria-independent apoptosis upon TNF-α treatment. To further determine the role of mitochondria in necrosis induced by TNF-α plus glucose deprivation, caspase-3 was expressed in MCF-7 cells to restore TNF-α-induced mitochondria-independent apoptosis (Figure 2E) (Scaffidi et al., 1998). In the presence of glucose, TNF-α induced the similar level of apoptosis in both MCF-7/vector and MCF-7/caspase-3 cells, and Bcl-xL over-expression completely blocked apoptosis in MCF-7/vector cells but did not affect apoptosis in MCF-7/caspase-3 cells (Figure 2F and S3D), indicating mitochondria-independent apoptosis in MCF-7/caspase-3 cells. In the absence of glucose, MCF-7/caspase-3 cells were less sensitive to necrosis and more susceptible to apoptosis upon TNF-α treatment compared to MCF-7/vector cells (Figure 2F), wherwas Bcl-xL over-expression completely suppressed necrosis but not apoptosis in MCF-7/caspase-3 cells induced by TNF-α plus glucose deprivation (Figure 2F). These observations suggest that mitochondrial injury plays a very important role in such a necrotic program. To test this hypothesis, we treated MCF-7 cells with carbonyl cyanide m-chlorophenyl hydrazone (CCCP) that causes an uncoupling of the mitochondrial proton gradient (Wallace and Starkov, 2000). CCCP induced mainly apoptosis in MCF-7 cells in the presence of glucose but necrosis in the absence of glucose (Figures S3E and S3F). Furthermore, ABT-737, a BH3 mimetic inhibitor of Bcl-2, Bcl-xL and Bcl-w (Oltersdorf et al., 2005), was also found to induce necrosis depending on glucose deprivation or glycolytic inhibition, and it showed the similar effects on MDA-MB-231 and Bcap37 cells (Figure S3G). By contrast, Bcl-xL/2 overexpression significantly prevented glucose deprivation-induced necrosis (Figure 2G). Taken together, our data show that mitochondrial impairment promotes necrosis induced by glucose deprivation.

### Metabolism has a decisive role in determining cell death

To further investigate what determines how TNF-α-treated MCF-7 cells die, we used a FBS-free, nutritionally deficient medium derived from DMEM. In the absence of nutrients, TNF-α predominantly induced necrotic cell death that was blocked by antimycin A and oligomycin A that suppressed the electron transfer chain and ATP synthase, but not by 2-Deoxy-D-glucose (2DG), an inhibitor of glycolysis (Figure 3A and S4A). Next, we checked the effect of nutrients on TNF-α-induced cell death by adding one nutrient each time to the nutrient-deficient medium. Screening with glucose, pyruvate, twenty amino acids and tricarboxylic acid cycle (TAC) intermediates showed that glucose, glycine and alanine almost completely shifted TNF-α-induced necrosis to apoptosis, while glutamine, serine and proline partially decreased necrosis and concomitantly increased apoptosis (Figures 3B –3G and S4A). With the addition of glucose, TNF-α-induced apoptosis was not inhibited by antimycin A or oligomycin A but switched by 2DG to necrosis that was further suppressed by the combination with mitochondrial inhibitors (Figure 3B). In contrast, with the addition of amino acids, TNF-α-induced cell death was almost completely blocked by mitochondrial inhibitors but not by 2DG (Figures 3C – 3G). These data suggest that it is the catabolism of these nutrients, not the nutrients themselves, which determines TNF-α-induced cell death in MCF-7 cells. This speculation was supported by additional results that some inhibitors, such as compound 968(Wang et al., 2010) and β-Chloro-L-alanine(Morino et al., 1979) that target catabolic pathway of glutamine and alanine, attenuated the effects afforded by glutamine and alanine (Figures S4B and S4C). By contrast, some TAC intermediates, such as citrate, α-ketoglutarate and malate, triggered necrosis alone or potentiated TNF-α-induced necrosis (Figures 3H and S5A), which was largely blocked by mitochondrial inhibitors (Figures 3H and S5B). These results suggest that mitochondrial catabolism of these TAC intermediates promotes glucose deprivation-induced necrosis.

**Figure 3.**
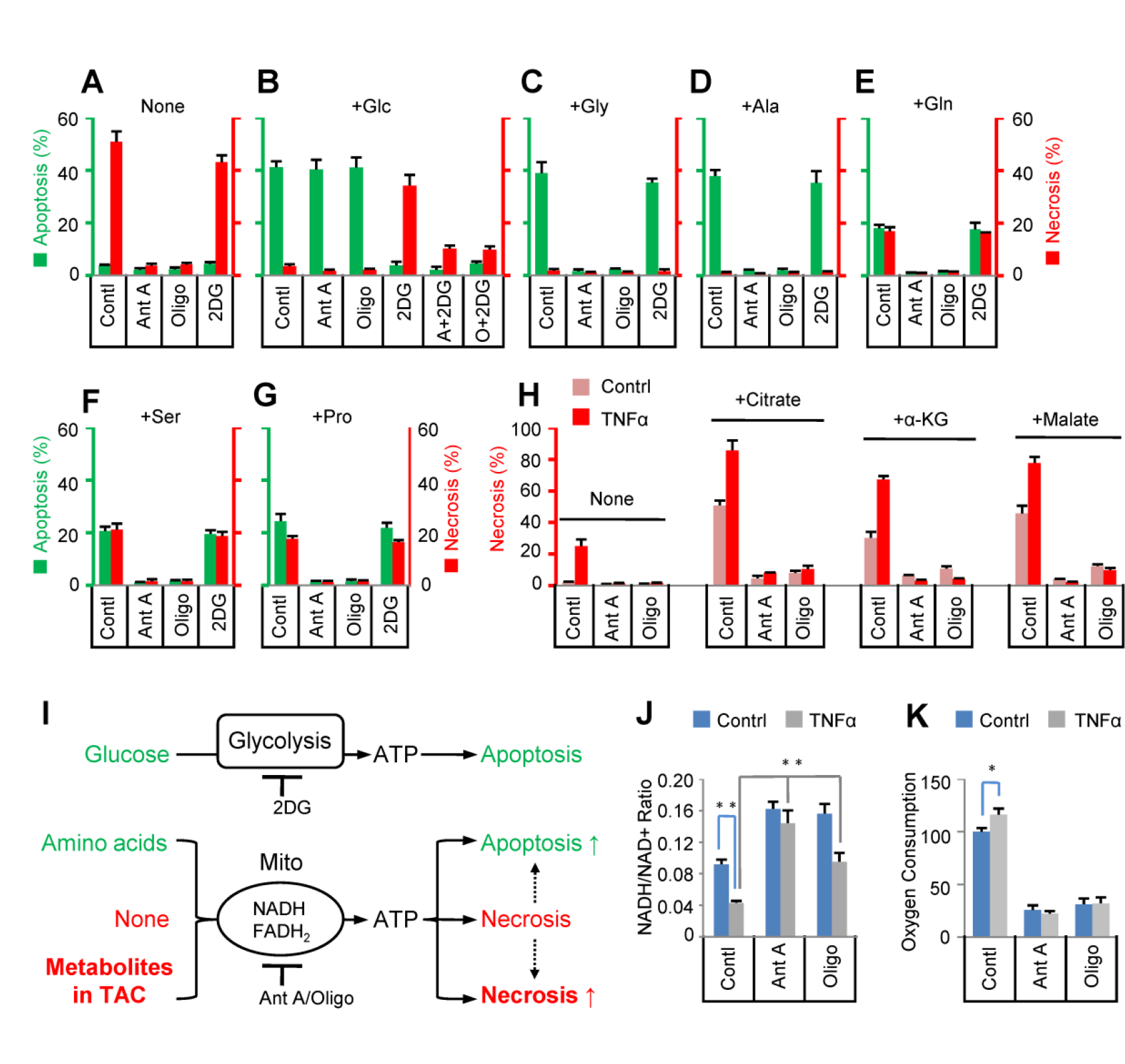
TNF-α-induced cell death type depends on metabolism. (A-G) The fraction of apoptotic and necrotic MCF-7/GC3AI cells in the nutrients-defined medium after treatment with 50 ng/ml TNF-α for 4h. The nutrient-deficient medium was supplied with nothing (A), 2 mM of glucose (B), 2 mM of glycine (C), 2 mM of alanine (D), 2 mM of glutamine (E), 2 mM of serine (F) or 2 mM of proline (G). Cells were pre-treated without or with antimycin A (1μM), oligomycin A (1μM) and/or 2DG (20 mM). Error bars indicate ± SD (n=3). (H) The fraction of necrotic MCF-7/GC3AI cells in the nutrients-defined medium after treatment with 50 ng/ml of TNF-α for 2h. The nutrient-deficient medium was supplied with nothing, 50 mM of citrate, 50 mM of α-ketoglutarate, or 50 mM of malate. Cells were pre-treated without or with antimycin A (1 μM), oligomycin A (1μM) and/or 2DG (20 mM). Error bars indicate ± SD (n=3). (I) Summary diagram of the relationship between metabolism and cell death. (J) The NADH/NAD^+^ ratio values of MCF-7/GC3AI cells cultured in the nutrient-deficient medium after treatment without or with 50 ng/ml of TNF-α for 2h. Cells were pre-treated without or with antimycin A (1 μM) or oligomycin A (1 μM) for 1 h. Error bars indicate ± SD (n=3). ** P<0.01 (t-test). (K) Oxygen consumption of MCF-7/GC3AI cells cultured in the nutrient-deficient medium after treatment with 50 ng/ml of TNF-α alone or in combination with antimycin A (1 μM) or oligomycin A (1 μM) for 1h. Data are represented as mean ± SD of triplicate wells. * P<0.05 (t-test). Experiments were repeated twice with similar results. See also Figure S4-S6.

The addition of nutrients to the nutrient-deficient medium showed no effect on ATP content in MCF-7 cells (Figure S4D). In the nutrient-deficient medium without or with the addition of amino acid, ATP generation in MCF-7 cells was completely blocked by mitochondrial inhibitors. In the presence of glucose, the ATP content of MCF-7 cells was not affected by mitochondrial inhibitors, but was totally eliminated by 2DG in combination with mitochondrial inhibitors (Figure S5D). Considering the effects of these treatments on TNF-α-induced cell death (Figures 3A-3G), these data support the idea that ATP generation is required for both TNF-α-induced apoptosis and necrosis. However, they did not explain what determined apoptotic or necrotic cell death.

Intracellular amino acids and TAC metabolites have to be converted to NADH/FADH_2_ to produce ATP through mitochondrial pathway (Figure 3I). Our results showed that the level of total amino acids, succinate and the NADH/NAD^+^ ratio in MCF-7 cells decreased with time in nutrient-deficient medium (Figures S6A-S6C), suggesting that the cells consumed these metabolites. Moreover, TNF-α gave rise to a further decrease in NADH/NAD^+^ ratio and an increase in oxygen consumption that were blocled by mitochondrial inhibitors (Figures 3J and 3K), indicating the promotion of mitochondrial metabolism by TNF-α. This most likely resulted from TNF-α-impaired mitochondria needing to consume more metabolites to maintain ATP level. Glycine, alanine, serine, proline and glutamine can be catabolized in mitochondria and concomitantly produce NADH or FADH_2_ (Figure S6D), and thus their addition to nutrient-deficient medium can maintain NADH/NAD^+^ ratio like the addition of glucose or TAC intermediates (Figure S6E). However, these additions exerted differential effects on TNF-α-induced cell death in MCF-7 cells and supported apoptosis and/or necrosis that were suppressed by mitochondrial and/or glycolytic inhibitors (Figure 3I). Taken all together, our data strongly suggest that how ATP is generated, i.e., ATP generation-driven catabolism, and not ATP or NADH/NAD^+^ itself, determines whether MCF-7 cells die and how cells die upon TNF-α treatment. This raises the possibility that the intracellular microenvironment set up by the catabolic conversion of substrates to products takes on a critical role in determining cell death.

### Metabolism-driven proton homeostasis plays a critical role in determining the type of cell death

The common participant shared by all these metabolic reactions are protons coupled with the consumption or generation of NADH + H^+^ and protonated carboxylic acid (Figure S6D). Glycolytic catabolism starting from glucose facilitated TNF-α-induced apoptosis (Figures 3B,3I, S1E and S1F). In the glycolytic pathway, glucose is split into pyruvic acids, producing net protons that can be used for mitochondrial ATP generation (Figure 4A). In contrast, in the absence of glucose, to generate ATP, mitochondria have to metabolized NADH + H^+^ or protonated TAC intermediates (polycarboxylic acids) and consume net protons(Figures 4A and 4B). These conditions led to or promoted TNF-α-induced necrosis(Figures 3A, 3H and 3I). The catabolism of some amino acids in mitochondria, coupled with the production of NADH + H^+^ and carboxylic acids (Figure S6D), could overall produce the net protons to support TNF-α-induced apoptosis (Figures 3C-3G and 3I).

**Figure 4.**
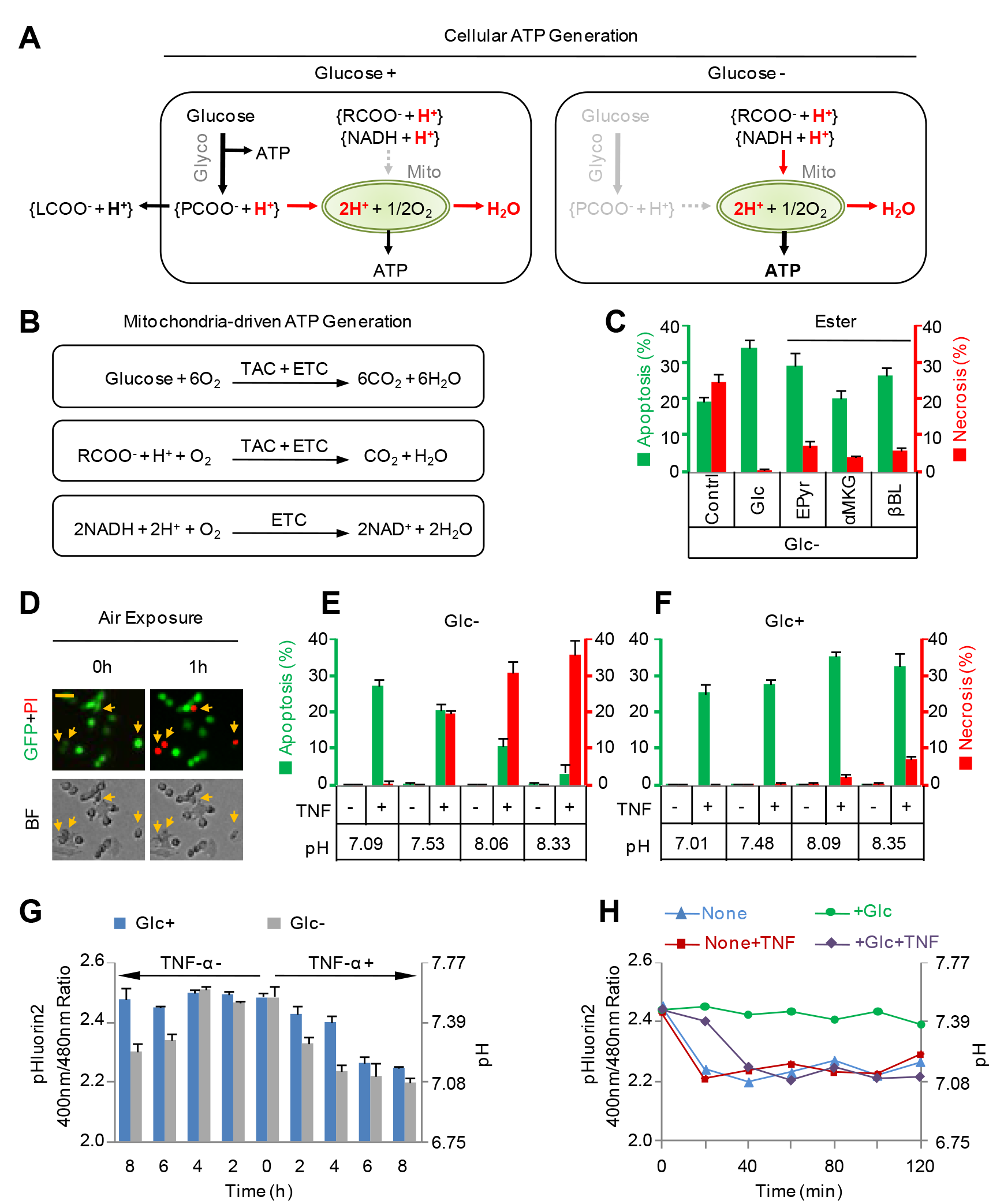
Proton homeostasis, but not the cytosolic pH, determined TNF-α-induced cell death type. (A) Protons-associated cellular ATP generation in the presence or absence of glucose. LCOOH, lactic acids; PCOOH, pyruvic acids; RCOOH, carboxylic acids, such as TAC intermediates and amino acids. (B) Diagram of the mitochondrial ATP generation pathway starting from glucose via TAC (tricarboxylic aicd cycle) and ETC (electron transfer chain), from TAC metabolites via TAC and ETC or from NADH + H+ through ETC. (C) The fraction of apoptotic and necrotic MCF-7/GC3AI cells after treatment with 50 ng/ml of TNF-α for 6h, and cells were cultured in the glucose-free medium supplied without or with 2 mM of glucose, 5mM of ethyl pyruvate (EPyr), 5 mM of dimethyl α-ketoglutarate (αMKG) or 5 mM of β-butyrolactone (βBL). Error bars indicate ± SD (n=3). (D) Fluorescent images of MCF-7/GC3AI cells in the complete medium exposed to air for 1h after they were treated with 50 ng/ml of TNF-α for 8h. Scale bar, 20 μm. (E and F) The fraction of apoptotic and necrotic MCF-7/GC3AI cells after treatment with 50 ng/ml of TNF-α for 6h in the glucose-free medium (E) or the complete medium (F) with different pH values. Error bars indicate ± SD (n=3). (G) The intracellular pH of MCF-7 cells after treatments without or with 50 ng/ml of TNF-α for different times as indicated in the presence or absence of glucose. Error bars indicate ± SD (n=3, 180 cells in total). (H) The intracellular pH of a single MCF-7 cell after treatment without or with 50 ng/ml of TNF-α in the nutrient-deficient medium supplied without or with 2mM of glucose. The result shown here is the representative one of 20 cells that finally underwent necrotic death. See also Figure S7 and S8.

To confirm the role of metabolism-driven proton homeostasis in TNF-α-induced cell death, we treated MCF-7 cells with carboxylic acid esters. If these esters can be rapidly hydrolyzed to carboxylic acids and alcohols in cells (Figure S7A), they will transiently produce intracellular protons. Among the esters we tested, including ethyl pyruvate, diethyl malate, dimethyl α-ketoglutarate, diethyl succinate and triethyl citrate, ethyl pyruvate and dimethyl α-ketoglutarate strongly protected cells against TNF-α-induced necrosis but not apoptosis in the absence of glucose (Figure 4C). In addition, a similar protective effect was also shown by β-butyrolactone, which can be hydrolyzed to β-hydroxybutyric acid (Figure 4C and S7B). In nutrient-deficient medium, ethyl pyruvate, dimethyl α-ketoglutarate and β-butyrolactone also reduced TNF-α-induced necrosis, while their unprotonated carboxylate counterparts did not (Figure S7C). These data strongly suggest that metabolism-driven proton homeostasis plays a critical role in determining TNF-α-induced cell death, and that proton generation promotes TNF-α-induced apoptosis while proton consumption results in TNF-α-induced necrosis.

### The cytosolic pH does not account for the determination of cell death type

When TNF-α-treated MCF-7 cells incubated in complete medium were exposed to air at room temperature, a shift of apoptosis in some cells to necrosis was observed (Figure 4D), similar to the phenomenon of cell death transition induced by TNF-α plus glucose deprivation (Figure S1B). Therefore, these effects may arise from a similar mechanism. Since exposure to room temperature by itself did not change the form of cell death (data not shown), we focused on the increased medium pH due to the absence of buffering by 5% CO_2_, which was also highly associated with the proton homeostasis.

We changed the pH of the medium by adding different concentrations of NaHCO_3_. Interestingly, we found that the acidic condition shifted TNF-α-induced necrosis to apoptosis while alkaline treatment supported necrosis in the absence of glucose (Figure 4E). To avoid the influence of NaHCO_3_ itself, we cultured MCF-7 cells in a CO_2_-free incubator with different media or buffer solutions that were adjusted to around pH 7.0, 7.5 or 8.0 using HCl and NaOH. Our results again demonstrated that the acidic environment promoted TNF-α-induced apoptosis while alkaline environment promoted TNF-α-induced necrosis in various conditions without glucose supplement (Figures S8A-S8C). Moreover, the medium pH did not affect the intracellular ATP level (Figure S8D). We used a pH-sensitive fluorescent protein, ratiometric pHluorin2 to monitor the intracellular pH (pH_i_). The excitation at 400 nm of pHluorin2 decreases with a corresponding increase in the excitation at 480 nm upon acidification(Mahon, 2011; Miesenbock et al., 1998). We confirmed that the medium or buffer solution indeed changed the pH_i_ in MCF-7 cells (Figure S8E). Intriguingly, in the presence of glucose, TNF-α still predominantly caused apoptosis and only slightly increased necrosis under alkaline conditions (Figures 4F and S8C). In the meantime, we also observed that the alkaline treatment did not change pH_i_ effectively in the presence of glucose (Figure S8E). This most likely resulted from the compensation by increased lactate production stimulated by alkaline conditions (Figure S8F).

The above results revealed that pH_i_ seems to be very important in TNF-α-induced cell death. We next checked whether TNF-α changed the intracellular pH_i_ of MCF-7 cells. When pHluorin2-expressing MCF-7 cells were treated with TNF-α, the ratio of fluorescence at 400/480 nm decreased in the presence of glucose, indicating intracellular acidification (Figure 4G). However, we also detected similar intracellular acidification, not the expected alkalinization, upon TNF-α treatment in combination with glucose deprivation. Glucose deprivation actually decreased the pH_i_ by itself after 6h (Figure 4G). Glucose-free medium still partially supports apoptosis, which may affect the pH_i_ measurement based on a population of cells. To avoid such interference, we measured the pH_i_ in single cells in nutrient-deficient medium where TNF-α primarily induced necrosis (Figures 3A and S4A). The pH_i_ quickly dropped to around pH 7.0 regardless of TNF-α treatment, and such a drop was inhibited or delayed by the addition of glucose (Figures 4H and S8G). These results suggest that TNF-α or glucose starvation triggers intracellular acidification. Considering the results obtained in the extracellular conditions with different pH values, it apparently is not the cytosolic pH that determines the form of cell death induced by TNF-α. Protons are not evenly distributed within cells, and some of them are compartmentalized by organelles whose differential proton concentration is maintained by metabolism. Therefore, altered metabolism may drive the redistribution of intracellular protons while the extracellular environment universally changes the pH in the cytoplasm including organelles. With this concept in place, we hypothesized that the cell death-associated lysosome, which contains a much higher concentration of protons than the cytosol, is centrally involved in regulating apoptosis and necrosis under these conditions.

### Severely lysosomal alkalinization is required for necrotic cell death

We next investigated the involvement of lysosomes in TNF-α-induced cell death. We use LysoTracker Blue, a blue fluorescent dye that stains acidic organelles, to stain lysosomes. We found that, in the presence of glucose, TNF-α did not decrease the fluorescent staining while glucose-deprivation by itself reduced fluorescent staining and the decrease was potentiated by TNF-α treatment (Figure 5A). Notably, in both conditions, apoptotic cells were well stained (Figure 5A), indicating that apoptosis did not affect the lysosomal acidification. To further confirm this observation, wild type MCF-7 cells were pre-stained with a LysoSensor Green dye that becomes more fluorescent in acidic environments and shows reduced or no fluorescent upon lysosomal alkalinization(Han and Burgess, 2010). Then we treated the pre-stained cells with TNF-α in the presence or absence of glucose. We observed similar results. TNF-α further decreased the fluorescence intensity already reduced by glucose deprivation (Figure S9A), suggesting that lysosomal alkalinization occurred. In nutrient-deficient medium, we similarly detected lysosomal alkalinization, and the addition of glucose, alanine or glycine, which supported TNF-α-induced apoptosis, suppressed lysosomal alkalinization (Figure S9C). In addition, we also observed that alkaline medium led to rapid deacidification of lysosomes in the absence of glucose, but only mildly alkalinized lysosomes in the presence of glucose. In contrast, acidic medium inhibited lysosomal alkalinization induced by glucose deprivation (Figure S9D). These results suggest that lysosomal alkalinization is closely associated with TNF-α-induced necrosis.

**Figure 5.**
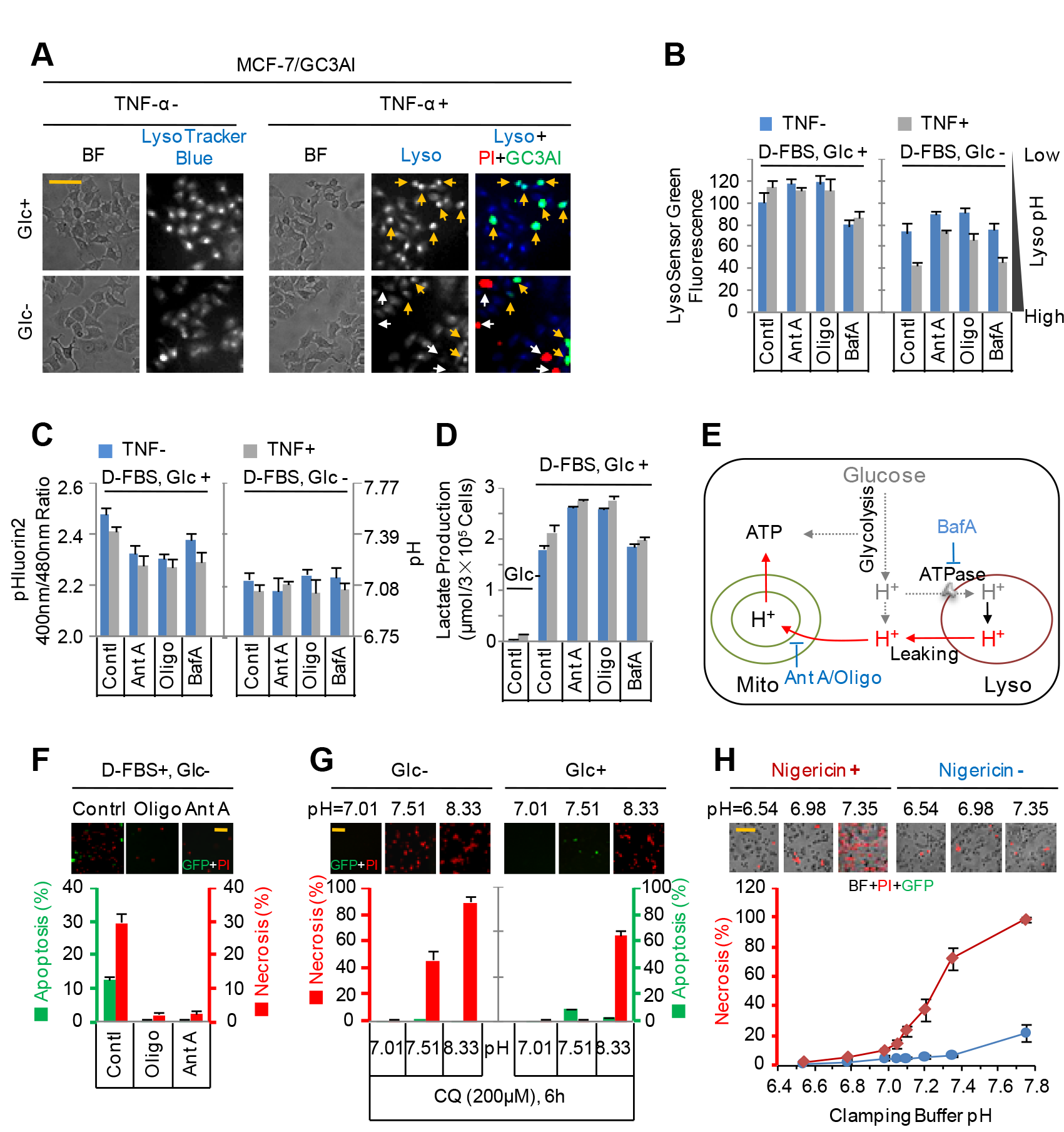
Proton homeostasis-driven lysosomal alkalinization determines the type of cell death. (A) Fluorescent images of MCF-7/GC3AI cells stained with LysoTracker Blue after treatments without or with 50 ng/ml of TNF-α for 5h in the presence or absence of glucose. Green fluorescence indicates apoptosis, red fluorescence shows necrosis, and blue fluorescence shows lysosomal acidification. (B) Lysosomal alkalinization of MCF-7/wt cells indicated by the pre-staining of LysoSensor Green after treatments without or with 50 ng/ml of TNF-α for 4h in the presence or absence of glucose. The medium supplemented with dialyzed FBS. Cells are pre-treated without or with antimycin A (1μM), oligomycin A (1μM) or bafilomycin A (1μM). The quantified fluorescence intensity was normalized by the number of cells. Error bars indicate ± SD (n=3, 180 cells in total). (C) The intracellular pH of MCF-7 cells measured by the stably expressed pHluorin2. Cells were treated in the conditions as described in (b). Error bars indicate ± SD (n=3, 180 cells in total). (D) Lactate production by MCF-7/wt cells after treatments as described in (b). Error bars indicate ± SD (n=4). (E) A model for the regulation of lysosomal acidification by metabolism-driven homeostasis. (F) Effect of mitochondrial inhibitors on cell death in MCF-7/GC3AI cells in the glucose-free medium with dialyzed FBS. Cells were pre-treated with 1 μM of oligomycin A or antimycin A before they were treated with 50 ng/ml of TNF-α for 6h. Error bars indicate ± SD (n=3). (G) The apoptotic and necrotic death in MCF-7/GC3AI cells after treatment with 200 μM of chloroquine for 6h in the complete medium or glucose-free medium with different pH values. Error bars indicate ± SD (n=3). (H) The necrotic death in MCF-7/GC3AI cells after incubation with the pH clamping buffers with different pH values for 8h. 25 μM of nigericin is used to equilibrate the pH outside and inside of cells. Error bars indicate ± SD (n=4). Scale bar in all panels, 40 μm. See also Figure S9 and S10.

In order to determine the effect of metabolism on lysosomal acidification, we cultured cells in medium containing dialyzed FBS. In the presence of glucose, antimycin A and oligomycin A slightly acidified both the cytosol and lysosomes(Figures 5B and5C) that possibly resulted from the increased glycolytic activity (Figure 5D). Glucose deprivation induced acidification of the cytosol while alkalinization of lysosomes (Figures 5B and 5C), suggesting proton release from the lysosome to the cytosol (Figure 5E). In the absence of glucose, TNF-α further exacerbated lysosomal alkalinization, an effect that was blocked by mitochondrial inhibitors (Figure 5B). Considering that mitochondrial inhibitors suppressed cell death induced by TNF-α plus glucose deprivation in this condition (Figure 5F), our data suggest that severe lysosomal alkalinization is most likely required for necrotic cell death induced by TNF-α plus glucose deprivation.

To further confirm the role of lysosomal alkalinization in necrosis, we treated MCF-7 cells with a lysosomotropic agent, chloroquine that can directly increase lysosomal pH(Klempner and Styrt, 1983). Our results showed that chloroquine did not significantly affect ATP content in the survived cells while induced significant necrosis in the absence of glucose but apoptosis in the presence of glucose (Figures 5G, S10A and S10B). However, in the alkaline medium, chloroquine induced necrosis regardless of glucose. In contrast, acidic medium blocked chloroquine-induced cell death (Figure 5G). Alkaline conditions potentiated lysosomal alkalinization directly (Figure S9D) and indirectly by deprotonating chloroquine to promote its accumulation in the lysosome (Solomon and Lee, 2009). Moreover, chloroquine synergistically promoted necrosis induced by TNF-α plus glucose deprivation (Figure S10C). Therefore, our data suggest that severe lysosomal alkalinization triggered necrosis.

In normal conditions where glucose metabolism produces protons to antagonize lysosomal alkalinization, chloroquine elicited obviously apoptosis alone by 24h of treatment besides potentiating TNF-α-induced apoptosis in MCF-7 cells (Figures 5G, S10B and S10C). This suggests that mild lysosomal alkalinization is sufficient by itself to induce apoptosis by a relative long time of treatment. Therefore, we tested the effect of bafilomycin A, an inhibitor of the H^+^-ATPase that maintains lysosomal acidification(Yoshimori et al., 1991). As expected, bafilomycin A mildly increased lysosomal pH while it decreased cytosolic pH (Figures 5C and 5D), and mainly resulted in apoptosis in MCF-7 cells after 48h of treatment (Figure S10D). We also observed that bafilomycin A-induced apoptosis was blocked by acid treatment while it was potentiated or shifted to necrosis in an alkaline environment (Figure S10E). Chloroquine and bafilomycin A also showed similar pH-dependent killing effects on MDA-MB-231 and Bcap37 cells (Figures S10F and S10G). In addition, long-term alkaline stress by itself induced apoptosis and/or necrosis in MCF-7, MDA-MB-231 and Bcap37 cells (Figures S10E and S10H). Therefore, it seems that the increasing lysosomal pH by itself determines whether cells undergo apoptosis or necrosis, which depends on the extent of lysosomal alkalinization. When we treated MCF-7 cells with pH clamping buffers, we observed that nigericin induced significant necrotic cell death in clamping buffer with a pH above 7.0 (Figure 5H). This suggests that pH 7.0 may be the lysosomal pH threshold that determines apoptosis or necrosis.

### Glucose is the major source to maintain proton homeostasis

In the presence of glucose, TNF-α did not impact lysosomal pH of MCF-7 cells treated with or without bafilomycin A (Figure 5B), thus it plus bafilomycin A did not shift apoptosis to necrosis (Figure S10C). Interestingly, in the absence of glucose, bafilomycin A had no effect on lysosomal pH, nor cell death (Figures 5C, 5D, S10I). This suggests that robust H+-ATPase activity requires the presence of glucose, consistent with previous reports (Kane, 1995; Sautin et al., 2005). Glucose-deprivation induced lysosomal alkalinization and led to cellular necrosis (Figures 6A and S11), an effect that was blocked by acidic medium and exacerbated by alkaline medium (Figures 6B, S12A and S12B). In contrast, in the presence of glucose, cells grew well in relatively alkaline conditions (Figure 6B), and this growth benefit should be provided by increased glycolytic activity (Figures S8F and S12C). Similar results were obtained with various cancer cell lines (Figure 6B). These results indicate that glucose not only maintains intracellular proton homeostasis but also protects cells against extracellular proton depletion. Among nutrients tested, including glucose, amino acids and pyruvate, we observed that only glucose deprivation induced cellular necrosis, although the withdrawal of some amino acids elicited apoptosis (Figure S12D), indicating a special role of glucose as the major proton source.

**Figure 6.**
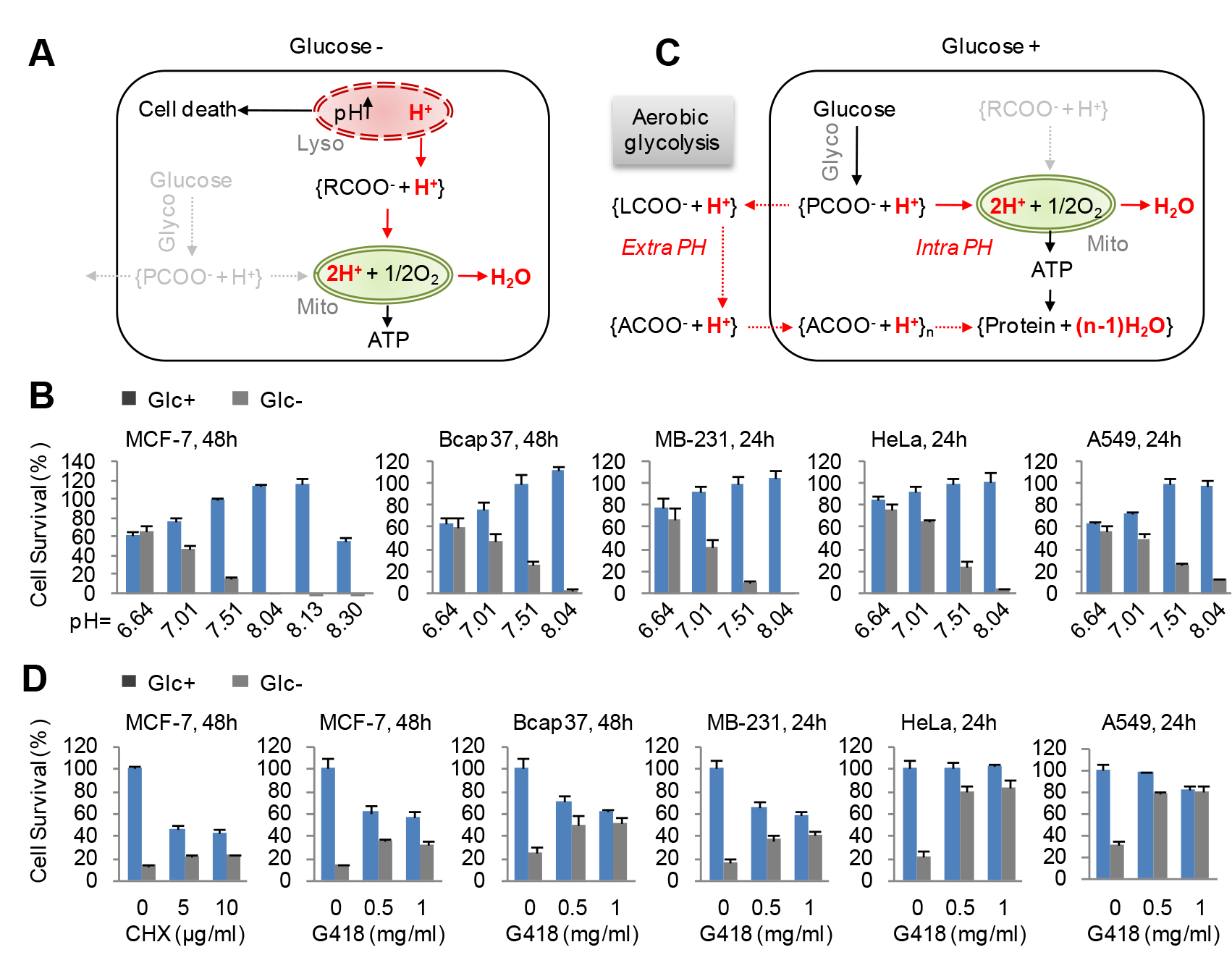
FGlucose is the major source in maintaining proton homeostasis. (A) A model for glucose deficiency-induced cell death mediated by alkalinized lysosomes. PCOOH, pyruvic acids; RCOOH, carboxylic acids. See Figure supplement 11 for details. (B) Survival of cells cultured in the complete medium or glucose-free medium with different pH values for 24h or 48h. Error bars indicate ± SD (n=3). (C) A model for the role of glucose in maintaining both intracellular and extracellular proton homeostasis (Intra PH and Extra PH). The Warburg Effect possibly compensates for the proton consumption by anabolism, in particular protein synthesis. PCOOH, pyruvic acids; RCOOH, carboxylic acids; LCOOH, lactic acids; ACOOH, amino acids. (D) Survival of cells cultured in the complete medium or glucose-free medium with protein synthesis inhibition treatments for 24h or 48h. Error bars indicate ± SD (n=3).

In addition to glucose, proliferating cells need to absorb a large quantity of amino acids for their active anabolism, in particular the synthesis of proteins (Hosios et al., 2016) (Figure 6C and 7A). The amino acids-associated protons are finally converted into waters during protein synthesis, which has to request a proton balancing compensation. This possibly specially is the case in vivo due to the limited extracellular space (Figure 7B). Such compensation can be optimally provided by lactic acid from aerobic glycolysis, the Warburg Effect (Figure 6C). Therefore, glucose-derived pyruvic acid can either enter mitochondria to maintain the ATP generation-involved intracellular proton homeostasis, or be converted into lactic acid to maintain the anabolism-associated extracellular proton homeostasis (Figure 6C). This speculation was further supported by the results that glucose deficiency-induced cell death was largely repressed by the inhibition of protein synthesis that consumes ATP and amino acids to drive both intracellular and extracellular proton consumption (Figure 6D).

**Figure 7.**
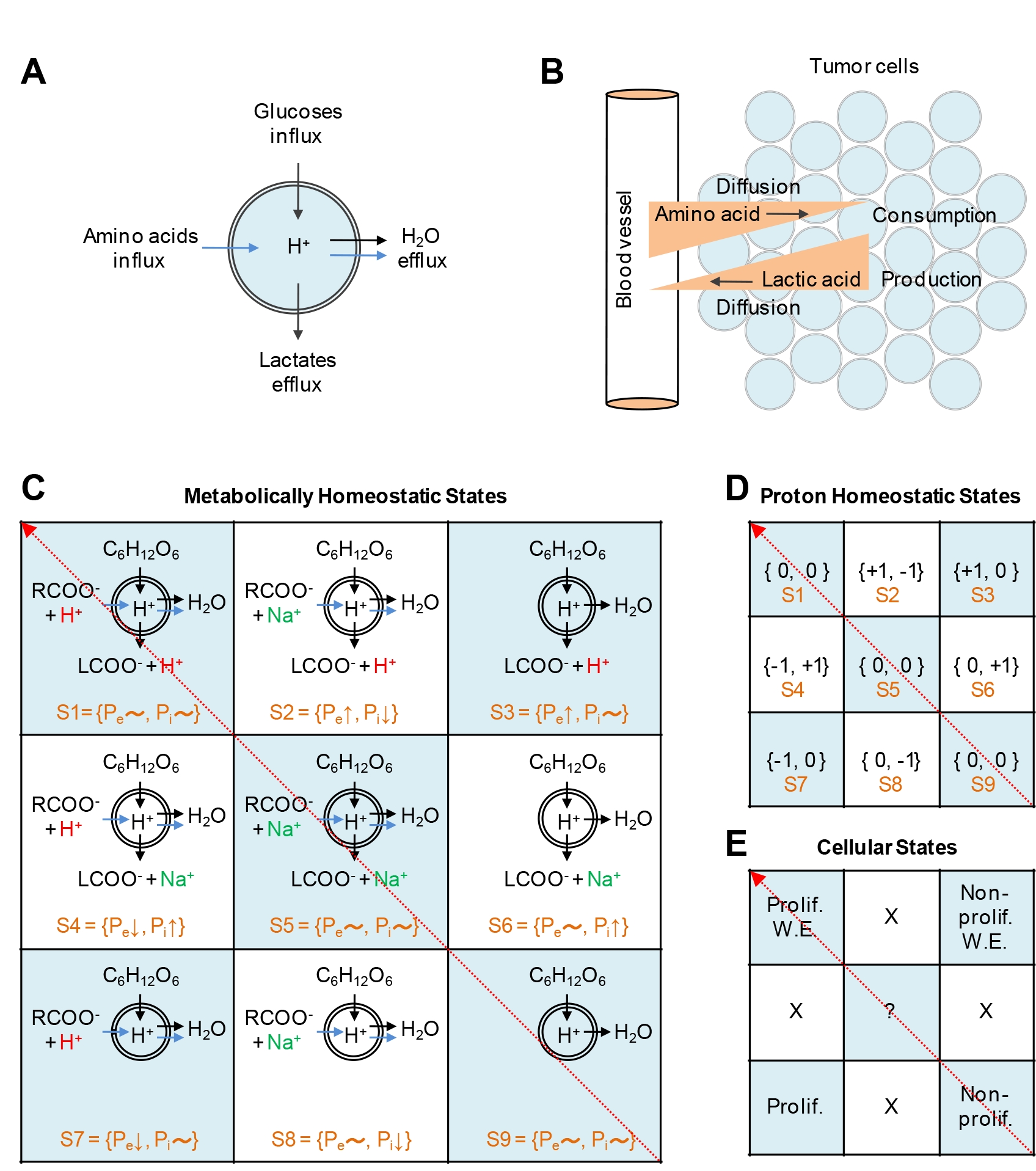
A theoretical model of cellular proton homeostasis. (A) A model for cellular metabolic flux. (B) A model showing the in vivo tumor microenvironment. The opposite gradient diffusions of amino acids and lactic acids between tumor cells and blood vessels compensate each other. (C) States of cellular metabolic homeostasis based on metabolic flux. RCOO^−^, carboxylates, mainly amino acids; Na+ represents cation. (D) States of cellular proton homeostasis based on metabolic states in (C). Pe, extracellular protons; P_i_, intracellular protons. +1, increased protons; −1, decreased protons; 0, balanced proton flux. (E) The states of cell proliferation corresponding to (C) and (D). S1, proliferation with the Warburg Effect; S3, non-proliferation with the Warburg Effect; S7, proliferation without the Warburg Effect; S9, non-proliferation. See also Figure S11-S13.

### The implicated role of the Warburg Effect

Proton homeostasis is one of the most important factors maintaining the microenvironment to enable metabolic reactions. Cells have several ways to maintain proton homeostasis mainly mediated by transporters (Parks et al., 2013), including antiporters, symporters and uniporters (Figure S13A). Proton transport is usually coupled with ions or metabolic groups, otherwise it will result in membrane potential. When coupled with ions, proton transport simultaneously perturbs the involved ion homeostasis. In contrast to the actual re-distribution of protons in other ways, the carboxylic metabolites-associated protons can be chemically expended or produced. In view of the obvious advantages, the metabolic pathway is naturally selected for cells to mainly maintain proton homeostasis during the evolution.

In the physiological condition, carboxylic groups usually exist as the unprotonated forms, thus cells could absorb or excrete protonated carboxylic acids or unprotonated carboxylates directly by single symporters or indirectly by symporters associated with antiporters (Figure S13A). In contrast, carboxylic metabolites are always produced or consumed as the protonated forms, thus the protonated carboxylic acids are simply the best participators in the cellular metabolic flux. Here we listed all nine possible metabolic states based on the order of protonated carboxylic acids, unprotonated carboxylates or nothing in the metabolic flux (Figure 7C). We analyzed in theory the intracellular or extracellular proton homeostasis based on the production or consumption of protons. If extracellular protons are absorbed or intracellular protons are consumed by cells, the extracellular proton concentration (P_e_) or intracellular proton concentration (P_i_) will decrease, and -1 is assigned to this situation. On the contrary, +1 is assigned to the increased protons that are excreted from or produced by cells. 0 is assigned to the balanced proton flux. Among all states of metabolic homeostasis in a 3X3 matrix, S2, S4, S6 and S8 cells are not able to maintain their intracellular proton homeostasis (Figures 7C and 7D), thus they are doomedly eliminated during the evolution. S1, S5 and S9 cells have the balanced proton homeostasis (Figure 7D). S1 represents proliferating cells with the Warburg Effect, including cancer cells, while S9 cells are corresponding to non-proliferating cells (Figure 7E). Up to date, no ion-coupled transporter responsible for excretion of metabolites is reported, which makes sure that lactic acid, not lactate, is carried out of cells by monocarboxylate transporters (Parks et al., 2013) to support proton homeostasis. This also means that S5 cells have to secrete carboxylates using symporters in combination with antiporters (Figure S13B). S5 having fussy metabolic flux might denote proliferating normal cells or a set of cells evolving to uncontrolled proliferating cancer cells (from S9 to S1). Interestingly, S3 and S7 cells could not keep alone their balanced extracellular environment but they do it together (Figure 7D). Upon co-survival with S3 cells, S7 cells can proliferate without the Warburg Effect that is actually provided by non-proliferating S3 cells (Figures 7D and 7E). The “so-called” reverse Warburg Effect was recently frequently reported in cancer-associated fibroblasts (Pavlides et al., 2009; Sotgia et al., 2012). These analyses indicate that maintaining proton homeostasis is a chemical requirement for proliferating cells.

## Discussion

Our experimental data indicate that glucose metabolism plays a critical role in maintaining cellular proton homeostasis, which supports cell survival or controls the choice of cell death. As illustrated in Figure S11, glycolysis and mitochondrial oxidative phosphorylation are the two major energy producing pathways in mammalian cells. In the glycolytic pathway, glucose molecules are split into pyruvates, concomitantly producing protons in the form of carboxylic acid and NADH + H^+^. In contrast, mitochondria consume protons by converting carboxylic acid and NADH + H^+^ to carbon dioxide and water (Figures 4A and 4B). These two pathways are normally integrated and cooperate with each other. In the absence of glucose, mitochondria continue to metabolize protons to generate ATP. Therefore, glucose deprivation eventually leads to the net consumption of protons, and meanwhile provokes compensatory lysosomal proton efflux that results in an increase in lysosomal pH and decrease in cytosolic pH. The resulting lysosomal alkalinization leads to apoptosis or necrosis depending on the extent of alkalinization. Mitochondrial impairments with apoptotic stimuli, such as TNF-α, CCCP and ABPT737, promote glucose deficiency-induced necrosis by accelerating the severe lysosomal alkalinization. In contrast, the lysosomotrpic treatment directly alkalinizes lysosomes and elicits necrosis in the absence of proton compensation from glycolysis, otherwise it may induce apoptosis upon the mild lysosomal alkalinization. Therefore, a major role of glycolysis (the proton producer) is to maintain proton homeostasis in cells together with lysosomes (the proton storer) and mitochondria (the proton consumer) to regulate cell survival.

As the proton storage bins, lysosomes can release protons to the cytosol and compensate the proton deficiency resulting from the absence of glucose. Such a compensation process may be critical to fast-growing cancer cells that often suffer from transient glucose starvation. However, a long-term glucose deficiency may over-alkalinize lysosomes to induced LCD. Considering that the pH-dependent enzymatic activity switch has been reported for lysosomal cathepsin A, a carboxypeptidase at acidic pH while deaminidase/esterase at neutral pH(Kolli and Garman, 2014), the neutralized luminal environment might change the specificity of some lysosomal hydrolases, and such enzymes may gain a new function to break down lysosomal membranes at neutral pH and thus trigger LCD.

In proliferating cells, in addition to mitochondrial ATP generation, the prevailing anabolism, in particular protein synthesis, also consumes a large number of protons (Figure 6C). Therefore, glucose metabolism needs to maintain both intracellular and extracellular proton homeostasis. Such a genetically metabolic reprogramming leads to aerobic glycolysis, the Warburg Effect (Vander Heiden et al., 2009). We experimentally and theoretically reveal the role of the Warburg Effect and explain the reverse Warburg Effect in cancer-associated fibroblasts that maintains the extracellular proton homeostasis for cancer cells. Our findings demonstrate the requirement of maintaining proton homeostasis for cell survival, and may expose a common Achilles’ heel of cancers, which might guide the design of general cocktail treatments for cancers. For example, Bcl-2 inhibitors in combination with glycolytic inhibition may synergistically kill cancers as observed in our current study (Figure S3G) and the study by others (Zagorodna et al., 2012).

## Experimental procedures

### Cell culture

MCF-7, MDA-MB-231, Bcap37, HeLa and A549 cells are obtained from ATCC. Stable cell lines were generated by lentivirus infection. Cells were maintained in high glucose DMEM supplemented with 10% fetal bovine serum (BioInd, Israel) and 50 IU penicillin/streptomycin (Invitrogen, USA) in a humidified atmosphere with 5% CO_2_ at 37°C. Experimental culture conditions were described in the Extended Experimental Procedures.

### Lysosomal acidification assessment

Lysosomal alkalinization was assessed by the staining with LysoTracker Blue (Invitrogen) or the pre-staining with LysoSensor Green (Invitrogen). See Supplemental Experimental Procedures for details.

### Lysosomal permeabilization assessment

Lysosomal permeabilization was assayed by a recently developed tool, EGFP-LGALS3 that binds permeabilized lysosomes and forms puncta (Aits et al., 2015). Cells expressing EGFP-LGALS3 were grown in twelve-well plates. After the desired treatments, cells were then imaged with an EVOS^^®^^ FLdigital inverted fluorescence microscope with 20× objective lens. The filter set for GFP imaging was Ex470/22, Em525/50.

### Cell death assay

Cell death was assayed by the fluorescence of cells stably expressing GC3AI in the medium containing PI. GFP-positive cells were counted as apoptosis, and cells only stained with PI were taken as necrotic cells. See Supplemental Experimental Procedures for details.

### Intracellular ATP assay

The intracellular ATP was measured using CellTiter-Glo Reagent (Promega) following the manufacturer’s instruction with minor modification. See Supplemental Experimental Procedures for details.

### Intracellular pH assay

The intracellular pH was supervised by a pH-sensitive fluorescent protein, pHluorin2 with a decrease in the excitation at 395 nm while a corresponding increase in the excitation at 475 nm upon acidification (Mahon, 2011; Miesenbock et al., 1998). See Supplemental Experimental Procedures for details.

### Lactate production assay

Lactate production was performed using the Lactate Assay kit (BioVision) according the manufacturer’s instruction. See Supplemental Experimental Procedures for details.

### Intracellular amino acids, succinate and NADH/NAD^+^ assay

The intracellular metabolites were measured using the Succinate Colorimetric Assay Kit, the L-Amino Acid Quantitation Colorimetric Kit and the NAD^+^/NADH Quantification Colorimetric Kit (BioVision) according to the user instructions. See Supplemental Experimental Procedures for details.

### Oxygen consumption assay

An XF24 Analyzer (Seahorse Biosciences, USA) was used to measure oxgen consumption in MCF-7 cells. See Supplemental Experimental Procedures for details.

### Statistics

Data are given as means ± SD. Statistical analyses were performed using unpaired, two-tailed Student’s t test for comparison between two groups. Asterisks in the figure indicated statistical significances (*, *p* < 0.05; **, *p* < 0.01).

## Authors Contributions

Conceptualization, B.L.; Investigation, Y.C., Y.W., P.X., and L.Q.; Formal Analysis, Y.C., Y.W. and B.L.; Writing-Original Draft, B.L.; Writing-Review & Editing, B.L. Y.C., and G.Y.; Funding Acquisition, B.L. and X.W.; Resources, Y.C., Y.W., P.X., L.Q. and M.L.; Supervision, B.L.

## Acknowledgements

We would like to thank Dr. John M. Kokontism, Dr. Wei Du (University of Chicago) and Dr. Guosheng Xiong (Agricultural Genomics Institute at Shenzhen China Academy of Agricultural Sciences, China) for constructive discussions and suggestions; Dr.Jiahuai Han (Xiamen University, China) for the caspase-3 construct. This work is supported by Grants 81372185 and 81472472 from Natural Science Foundation of China, and Grant TD12-5025 from Tianjin Municipal Innovation Program.

